# In vivo phosphoproteomics reveals pathogenic signaling changes in diabetic islets

**DOI:** 10.1101/439638

**Authors:** Francesca Sacco, Anett Seelig, Sean J. Humphrey, Natalie Krahmer, Francesco Volta, Alessio Reggio, Jantje Gerdes, Matthias Mann

**Affiliations:** Proteomics and Signal Transduction, Max-Planck Institute of Biochemistry, 82152 Martinsried, Germany; Helmholtz Diabetes Center (HMGU) and German Center for Diabetes Research (DZD), 85748 Garching, Munich, Germany; School of Life and Environmental Sciences, The University of Sydney, Sydney NSW 2006, Australia; Department of Biology, University of Rome Tor Vergata, 00100 Rome, Italy

## Abstract

Progressive decline of pancreatic beta cells function is key to the pathogenesis of type 2 diabetes. Protein phosphorylation is the central mechanism controlling glucose-stimulated insulin secretion in beta cells. However, if and how signaling networks are remodeled in diabetic islets *in vivo* remain unknowns. Here we applied high-sensitivity mass spectrometry-based proteomics and quantified the levels of about 6,500 proteins and 13,000 phosphopeptides in islets of obese diabetic mice and matched controls. This highlighted drastic remodeling of key kinase hubs and signaling pathways. We integrated our phosphoproteomic dataset with a literature-derived signaling network, which revealed a crucial and conserved role of GSK3 kinase in the control of the beta cells-specific transcription factor PDX1 and insulin secretion, which we functionally verified. Our resource will enable the community to investigate potential mechanisms and drug targets in type 2 diabetes.

## INTRODUCTION

The metabolic syndrome is caused by a complex interplay between genetic, epigenetic and lifestyle factors, including physical activity and diet (Zimmet et al., 2001). In particular, type 2 diabetes (T2D) mellitus is a major public health issue, accompanied by a chronic decline of glycemic control. The pathophysiology of type 2 diabetes starts with insulin resistance and proceeds to beta cell dysfunction (Prasad and Groop, 2015). Growing evidence implicates beta cells dysfunction as a key event during T2D development (Saisho, 2015). Genome-wide association (GWAS) studies have identified multiple risk variants for T2D with a primary role in beta cell function, highlighting their importance in the development of T2D (McCarthy and Hattersley, 2008). However, our knowledge of the molecular mechanisms underlying the development of beta cell dysfunction is still far from complete. A systems-wide understanding of beta cell dysregulation obtained by unbiased methods could contribute to the treatment and prevention of this disease. Large-scale “omics” technologies, particularly transcriptomics but also mass spectrometry (MS)-based proteomics, have been applied to characterize islets isolated from different T2D animal models and human cadavers (El Ouaamari et al., 2015; Hou et al., 2017; Lu et al., 2008; Segerstolpe et al., 2016). While these analyses provided sets of differentially expressed genes and proteins involved in beta cell failure, by their design they did not capture the crucial signaling alterations at the level of phosphorylation networks. As phosphorylation is a major regulator of insulin secretion and can also be modulated pharmacologically, its global investigation would be highly desirable.

Phosphorylation can be measured at a large scale by proteomics methods, but this involves specific enrichment for phosphorylated peptides and typically requires hundred-fold larger input material than proteomics measurements. Consequently, a major obstacle to the global characterization of changes in T2D islets by phosphoproteomics has been the extremely limited amount of material that can be extracted from pancreatic islets. Our group has recently described a MS-based phosphoproteomics workflow, termed ‘EasyPhos’, which enables streamlined and large-scale phosphoproteome analysis over multiple experimental conditions (Humphrey et al., 2015). Very recently, we have made further developments to this method, improving its sensitivity several-fold (Humphrey et al., 2018). We reasoned that this workflow might enable the in-depth characterization of changes in signaling networks of islets isolated from diabetic compared to control mice. For this purpose, we employed mice that carry an autosomal recessive mutation in the leptin receptor, the standard model in the field. These Lepr^db/db^ mice become obese from 3–4 weeks of age, and develop hyperglycamia as early as 4–8 weeks (Chen et al., 1996). To obtain a comprehensive view of global signaling network rewiring, we combined the phosphoproteomic analysis of diabetic islets with an in-depth characterization of the proteome changes. Our analysis revealed a drastic remodeling of the proteome and phosphoproteome, including numerous proteins implicated in insulin secretion control, glucose uptake and metabolism. To functionally interpret our datasets we applied a recently developed bioinformatics workflow (Sacco et al., 2016b). This highlighted a novel GSK3-dependent signaling axis contributing to beta cell failure, whose molecular mechanism we investigated in mouse and rat cells as well as in human islet models.

## RESULTS

### In-depth proteomic and phosphoproteomic characterization of db/db islets

To obtain a comprehensive perspective of the mechanisms underlying the insulin secretion failure occurring in diabetic islets, we measured the proteome and phosphoproteome of pancreatic islets isolated from 13 week old homozygous C57BLKS-Lepr^db^ mice. As expected, these “db/db” mice were obese and hyperglycaemic (**Fig. S1A-B**). As negative controls, we used age-matched C57BLKS-Lepr^db^/+, because these heterozygous mice have the same genetic background (hereafter referred to as “Ctrl”) (**Fig. 1A**).

**Figure 1.**
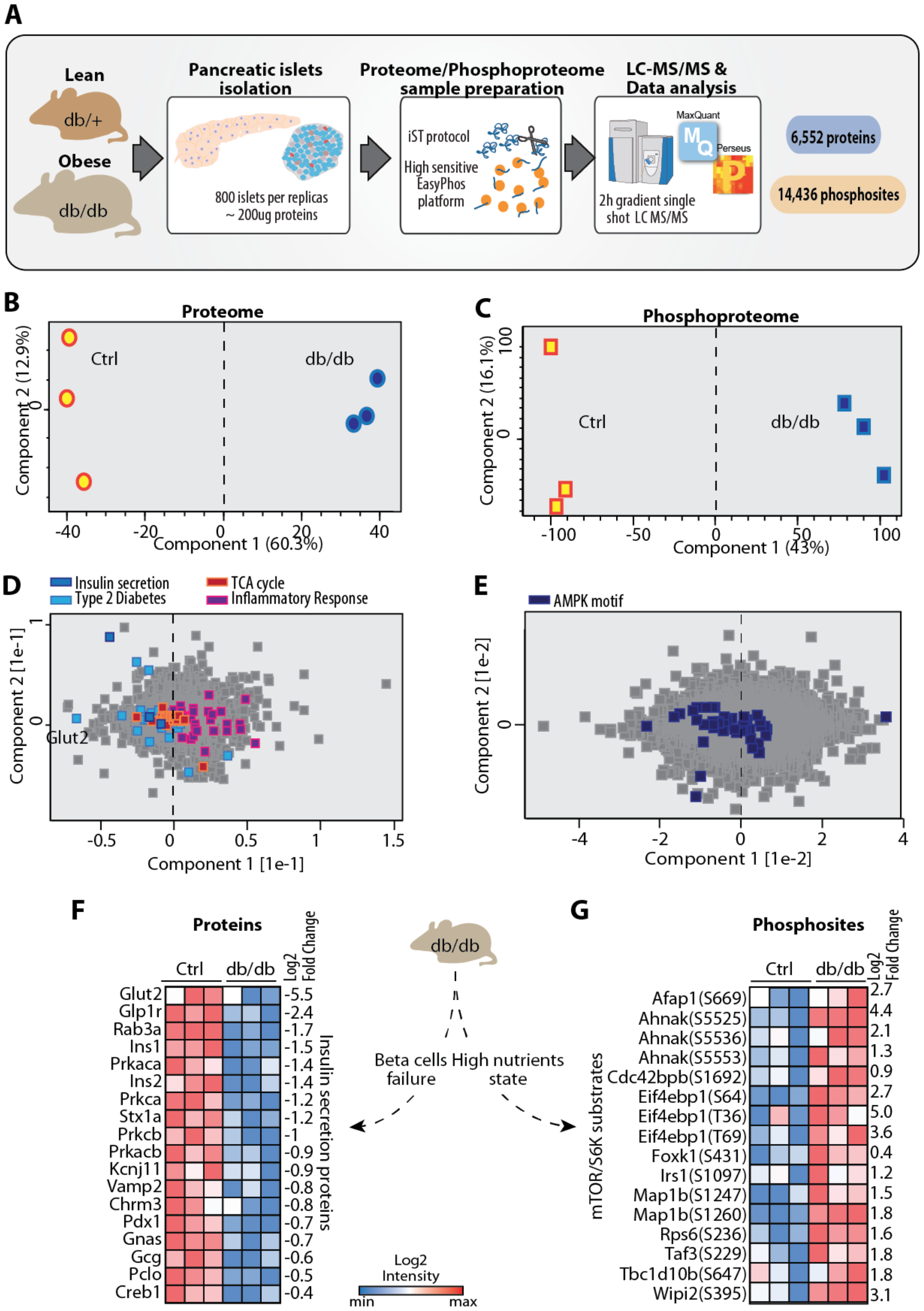
Our experimental strategy. (**A**) Experimental workflow to analyze the proteome and phosphoproteom of pancreatic islets derived from diabetic (db/db) and control (db/+) mice. Principal component analysis (PCA) of proteom (**B**) and phosphoproteom (**C**) data. The loadings of the PCA in B and C reveal that the proteins (**D**) and phosphosites (**E**) responsible for driving the segregation in component 1 are significantly enriched in the GO BPs and Kegg pathways and kinase substrate motifs indicated (FDR<0.07). **F**) Heat map of protein levels of the labeled insulin secretion-associated proteins in control and db/db islets upon drug and glucose treatments. (**G**) Heat map of phosphorylation levels of the indicated phosphosites in control and db/db islets upon drug and glucose treatments.

We applied a label-free quantification approach followed by high-resolution LC-MS performed in a single-run format on the newest generation Q Exactive mass spectrometers (Kelstrup et al., 2018; Kulak et al., 2014). Combined with the latest developments of our EasyPhos workflow, this allowed us to work on single mice for the proteome, whereas we harvested islets from three mice for the phosphoproteome measurements, yielding 200 μg of islet proteins. This experimental strategy enabled us to reliably quantify 6,500 proteins (**Fig. 1A**, **Supplementary Table S1**) and more than 14,000 different phosphorylation events mapping to 3,821 proteins (**Table S2; Fig. S2A-B**). In total 88% of the phosphoproteome were localized with high confidence to a single amino acid sequence location (**Fig. S2C**), with a median localization probability greater than 0.99 (**Experimental Methods**). These are excellent values, especially when considering the minimal amount of in vivo material, and the fact that no exogenous stimulation was employed.

Both proteome and phosphoproteome measurements where highly accurate and reproducible, with Pearson correlation coefficients of biological replicates ranging between 0.85 (lowest for phosphoproteome measurements) and 0.99 (highest for proteome measurements) (**Fig. S2D-E**). In agreement with other MS-based studies (Sharma et al., 2014), the majority of phosphorylation events in pancreatic islets occurred on serine residues (88%), followed by threonine (11%), whereas phosphotyrosines accounted for less than 1% of quantified phosphosites. Compared to the community database PhosphoSitePlus (Hornbeck et al., 2015), 87% of our quantified phosphosites were identical — a reassuring proportion considering the small sample amount and the depth of the analysis (**Figure S3A-B**). Only 268 out of these 10,994 sites are annotated as ‘regulatory’, meaning that the functional role or upstream cognate kinases of the vast majority of phosphorylation events quantified here have not been investigated to date.

Principal component analysis (PCA) of both our proteomic and phosphoproteomic data clearly classified pancreatic islets according to their diabetic status (**Fig. 1B-C**). AMPK inhibition, mTOR hyperactivation and insulin secretion failure are features of type 2 diabetic islets (Blandino-Rosano et al., 2012; Kahn et al., 2006). The drivers of the discrimination (component 1 of the PCA loadings) at the proteome and phosphoproteome level are enriched in annotations with these processes, providing a positive control for our proteomics measurements (**Fig. 1D-E**). Likewise, db/db-upregulated proteins were also significantly enriched for inflammation processes, which is known to correlate with obesity and T2D progression (Donath et al., 2013). Additionally, key proteins involved in insulin secretion were significantly down-regulated in db/db islets with fold-changes between 0.5 and five-fold whereas the phosphorylation of almost all the mTOR/S6K substrates was increased by similar amounts (**FDR < 0.05**; **Fig. 1F-G**). We conclude that our proteomic screen recapitulated known biology of diabetic islets - in particular with regards to the effects of chronic high nutrients on mTOR hyperactivaton and insulin secretion failure - while generating a vast resource of previously unknown, regulated events.

We next compared our dataset to previous proteomic studies characterizing proteome remodeling in pancreatic islets of murine and rat T2D models. In insulin-resistant MKR mice, only 159 regulated proteins were reported (Lu et al., 2008), however, 45% of them were also regulated in our much deeper dataset (**Fig. S4A**). Likewise, in a study of ob/ob and high-fat diet mice (El Ouaamari et al., 2015) 198 of the 272 significantly regulated proteins were likewise in our set. In addition to highly significant overlap of identifications, the Pearson correlation coefficients to our study were also relatively high (from 0.7 to 0.83) (**Fig. S4B** and **Fig. S5A-D**). In contrast, the 2,372 regulated proteins in non-obese GK rats (Hou et al., 2017) had less, although still statistically significant overlap with our data (35%; p < 0.05). We attribute this to the fact that the previous models all involve obesity and insulin resistance, whereas beta-cell failure is apparently quite different in the GK model. Interestingly, a large fraction of the 107 genes significantly modulated in islets derived from all four different T2D models were involved in the ER stress response (FDR < 0.05; **Fig. S4C and S5E**). Glucolipotoxicity triggers the accumulation of misfolded proteins that are removed from the ER and subjected to endoplasmic reticulum-associated degradation (ERAD). Misfolded proteins are translocated to the cytosol, ubiquitylated and degraded in the proteasome (Karunakaran et al., 2012). Thus, these unbiased proteomic screens suggest that the upregulation of different components of the ubiquitin/proteasome system is a major common adaptation mechanism of type 2 diabetic islets to glucolipotoxicity caused by the accumulation of misfolded proteins.

GWAS studies have identified several DNA loci associated with T2D (Scott et al., 2017), however, whether and how these genetic variants lead to changes in gene expression and are functionally connected to T2D generally remains unclear. We observed a significant overlap between genes carrying SNPs in T2D patients and db/db modulated proteins: more than 60% of the GWAS genes quantified at the protein level by our screen were significantly modulated (**Fig. S4D**). This suggests that protein levels might in some cases be linked to DNA variants of the respective gene products by affecting gene expression, protein stability, or degradation. This connection also implies that some of the proteins differentially expressed in the islets of this mouse model of T2D may also be affected in islets of T2D patients.

### Drastic remodeling of the proteome and phosphoproteome in db/db islets

We next investigated to what extent the proteome and phosphoproteome are altered in dysfunctional islets in our db/db mice. Around 35% of the proteome were modulated in db/db islets at an FDR < 0.07 (**Fig. S6A, S7A** and **Table S1**) and more than 25% of phosphopeptides on over 40% of phosphoproteins (**Fig. S6B, S7B-C** and **Table S2**). Interestingly, the levels of 72% of these phosphopeptides are increased in db/db with respect to control islets. We interpret this observation to reflect a more active global signaling due to the high nutrition state.

Combined analysis of the phosphoproteome and proteome data showed that phosphorylation sites were evenly detected, irrespective of protein abundance and that for 82% of the quantified phosphorylation sites, the corresponding protein was also quantified (**Fig. S3C, D, S7D**). When normalizing for the protein levels, more than 60% of sites were still significantly regulated, while the regulation of the remaining proteins was due to a combination of changing phosphorylation and protein levels. In 65% of those case, phosphorylation and protein levels both increased (46%) or decreased (18%) (**Fig. S7E**). Together, our data reveals that a full third of the proteome and a quarter of the phosphoproteome are changed in our diabetes model. This result also highlights the need for joint analysis of protein and phosphorylation changes for a complete view of the molecular changes underlying pancreatic islet dysfunction.

We next asked which biological processes were most altered in db/db islets and at which molecular level. For the proteome, a strong down-regulation of proteins involved in insulin secretion (at least two-fold and up to 100-fold) correlated with an impairment of mitochondrial metabolic processes, such as TCA cycle and OXPHOS, and with an increased lipid metabolism (**Fig. S7F**). Down-regulation of annotated insulin secretion proteins was also associated with increased ER and oxidative stress. Interestingly, db/db islets showed a positive enrichment of proteins involved in cell-cell adhesion and cell migration. At the signaling level, the mTOR pathway and autophagy, a down-stream process, were positively enriched in db/db islets. These changes occurred only at the phosphoproteome level and will be discussed in more detail below.

### Key signaling pathways are remodeled in diabetic islets

To investigate the mechanisms responsible for the phosphoproteome level changes, we extracted the kinase substrate motifs (Keshava Prasad et al., 2009) significantly changed in db/db islets (**Fig. 2A**) (Fisher exact test, FDR < 0.05; see Experimental Procedures). Consistent with high nutrient-induced mTOR hyperactivation, the substrate motif of p70S6K was enriched, while the AMPK motif was down-regulated. Other key kinase motifs, such as those of ERK1/2 and AKT, were also down-regulated. Additionally, the substrate motifs of key kinases previously implicated in the regulation of insulin secretion, including PKA, PKC and CaMK (Sacco et al., 2016a; Shibasaki et al., 2007), were down-regulated, further explaining the inability of diabetic beta cells to properly secrete insulin after glucose stimulation. Enrichment or depletion of a specific substrate motif may be the consequence of the modulation of kinase activity or concentration and the proteome levels can help to unravel this question. We observed all of the catalytic subunits of PKA and PKC to be consistently down-regulated in db/db islets, as well as the MAPKs ERK1 and ERK2 (**Fig. 2B**).

**Figure 2.**
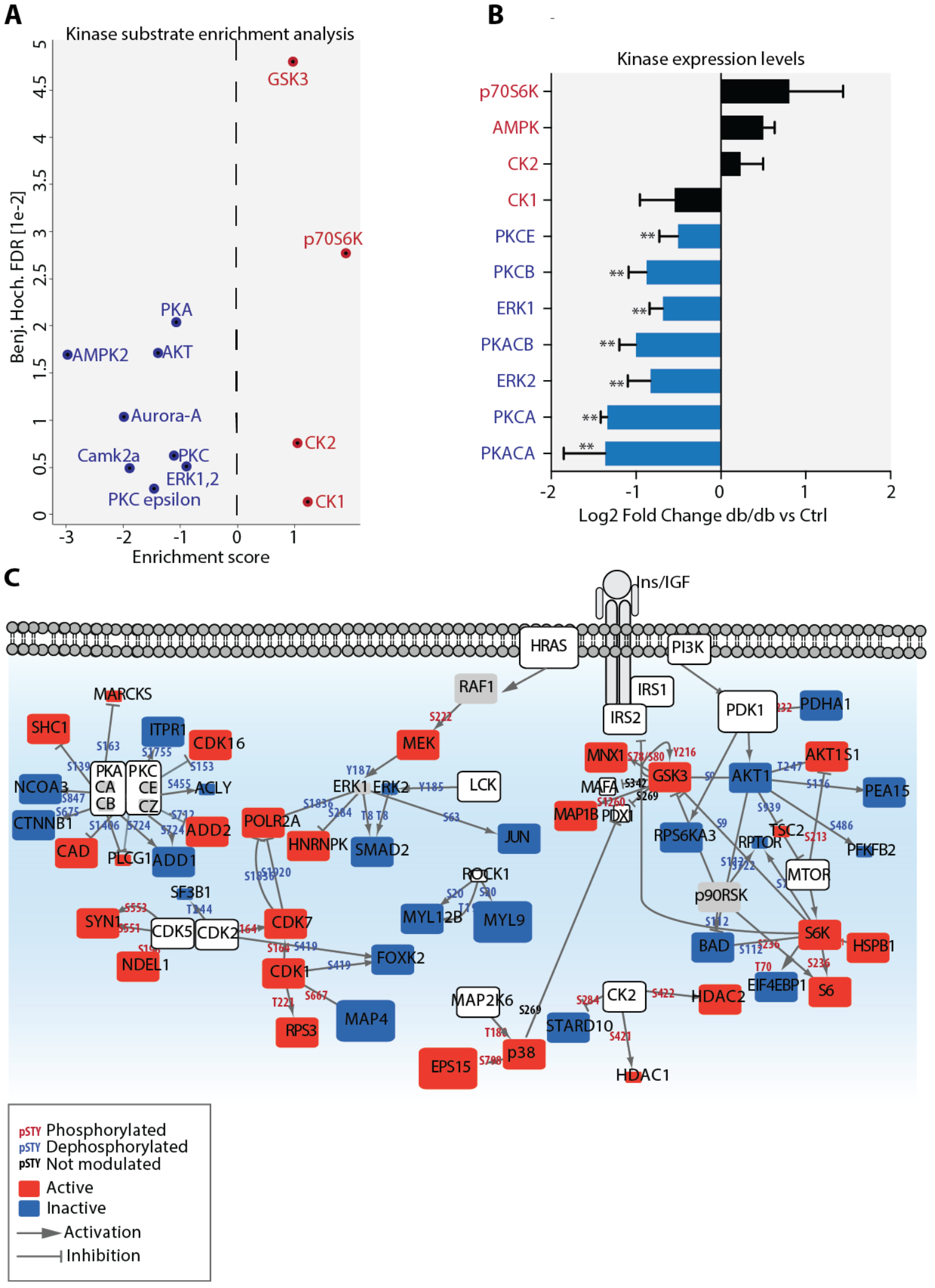
Key signaling pathways are modulated in db/db islets. (**A**) Kinase substrates motifs significantly overrepresented in significantly modulated phosphosites in db/db islets (Benjamin Hochberg FDR < 0.05). **B**) Log2 fold change of the expression levels of the indicated kinases in db/db islets with respect to control islets. **C**) The phosphoproteome and proteome changes observed in db/db islets were mapped onto a literature curated signaling network. Node size is proportional to the protein expression change in db/db islets. Blue and red nodes are inactivated or activated in db/db islets, respectively. Phosphorylation and dephosphorylation reactions on specific residues are represented as edges between nodes. Phosphosites are colored according to their phosphorylation state in db/db islets, as indicated in the legend.

As our data clearly indicate that key kinases are modulated in db/db islets, we next wanted to understand how signaling pathways are globally rewired in conditions of beta cells failure. We previously developed an approach that integrates large-scale quantitative phosphoproteomic and proteomic data with a literature-derived, proteome-wide signaling network (Sacco et al., 2016b). Exploration of the activation/inactivation pattern in the resulting graph confirmed and extended our previous observations, highlighting that all PKA and PKC substrates in our dataset were significantly dephosphorylated in the db/db islets, and that the expression levels of the PKA and PKC catalytic subunits was also decreased (**Fig. 2C**). Several G protein-coupled receptors, including Gpr119, Glp1 and Gpr40, have attracted considerable interest as T2D drug targets, due to their potential positive effects on insulin secretion *via* indirect activation of PKA and PKC (Bailey et al., 2016). Our data indicate that PKA and PKC are down-regulated, strongly suggesting that drug efficacy considerations should take the results of proteomics measurements into account.

Interestingly, we found that phosphorylation of the activating residue Thr8 of the SMAD2 transcription factor, which leads to its inactivation (Funaba et al., 2002), was significantly decreased. Consistently, Smad2βKO mice have striking islet hyperplasia together with defective glucose-responsive insulin secretion (Nomura et al., 2014).

The high-nutrient–induced mTOR hyperactivation that we observed in diabetic islets leads to inhibition of AKT through the well-characterized p70S6K-IRS1/2 negative feedback loop (Hsu et al., 2011). Decreased AKT activity in turn decreases phosphorylation of its direct substrate, the inhibitory Ser9 of GSK3, which we indeed observed in our dataset. Likewise, phosphorylation of the activating Tyr216 of GSK3 was significantly increased, further confirming increased GSK3 activity in db/db islets. Ser269 of the key beta cell specific transcription factor PDX1 is a direct substrate of GSK3, and phosphorylation of this residue causes its proteasomal degradation (Humphrey et al., 2010). In db/db islets the phosphorylation of the Ser269 of PDX1 was increased, while the protein level of PDX1 impaired (**Fig. S8**). Together, our joined literature and proteomic network, suggests a signaling axis starting from mTOR hyperactivation, GSK3 activation and PDX1 degradation, whose functional role in diabetic islets we investigate below.

### PDX1 suppression down-regulates GLUT2 and inhibits glycolysis in db/db islets

Given our finding that GSK3 is hyperactivated in diabetic islets, leading to PDX1 degradation and the established importance of PDX1 in beta cell physiology, we asked whether the expression of downstream targets of PDX1 was also impaired in beta cells. Such targets include the glycolytic enzyme glucokinase, the insulin genes, the major murine islet glucose transporter GLUT2, and the two transcription factors NKX6.1 and HNF4A (Gao et al., 2014). Remarkably, all of these PDX1 targets were significantly down-regulated in db/db islets (except for HNF4A, which we did not identify), with GLUT2 protein levels drastically decreasing by 64-fold in db/db islets compared to control islets (**Fig. 3A**). In rodents, GLUT2 is the only glucose transporter detected in normal pancreatic islets and therefore it has a crucial role in the insulin secretion pathway. In the absence of GLUT2, there is no compensatory expression of either GLUT1 and GLUT3 in islets (Guillam et al., 2000), and accordingly, our in-depth proteomic analysis detected neither GLUT1 nor GLUT3. Decreased expression of GLUT2 strongly impairs glucose ability to effect insulin biosynthesis and secretion and this correlates with a strong impairment of whole body glucose utilization (Guillam et al., 2000). Consistently, our combined proteomic and literature mining approach revealed that in db/db islets most of the key enzymes involved in glycolysis and the TCA cycle were significantly down-regulated, globally impairing glucose metabolism (**Fig. 3B**). In particular, we observed that the expression levels of almost all the subunits of the respiratory enzymes were also significantly decreased – by two-fold on average. Upon increased plasma glucose concentrations, this will impair a corresponding rise of ATP and the lower [ATP]/[ADP] concentration should negatively affect downstream components of the insulin secretion machinery. Indeed, the proteomics data indicated a significant downregulation of the protein level of the K_ATP_ channel subunits (1.5-fold, **Table S1**). Thus our data provide a molecular explanation for the insulin secretion failure observed in islets from 10-13 weeks old db/db mice (Do et al., 2014; Guan et al., 2016; Kim et al., 2015; Kondo et al., 2012).

**Figure 3.**
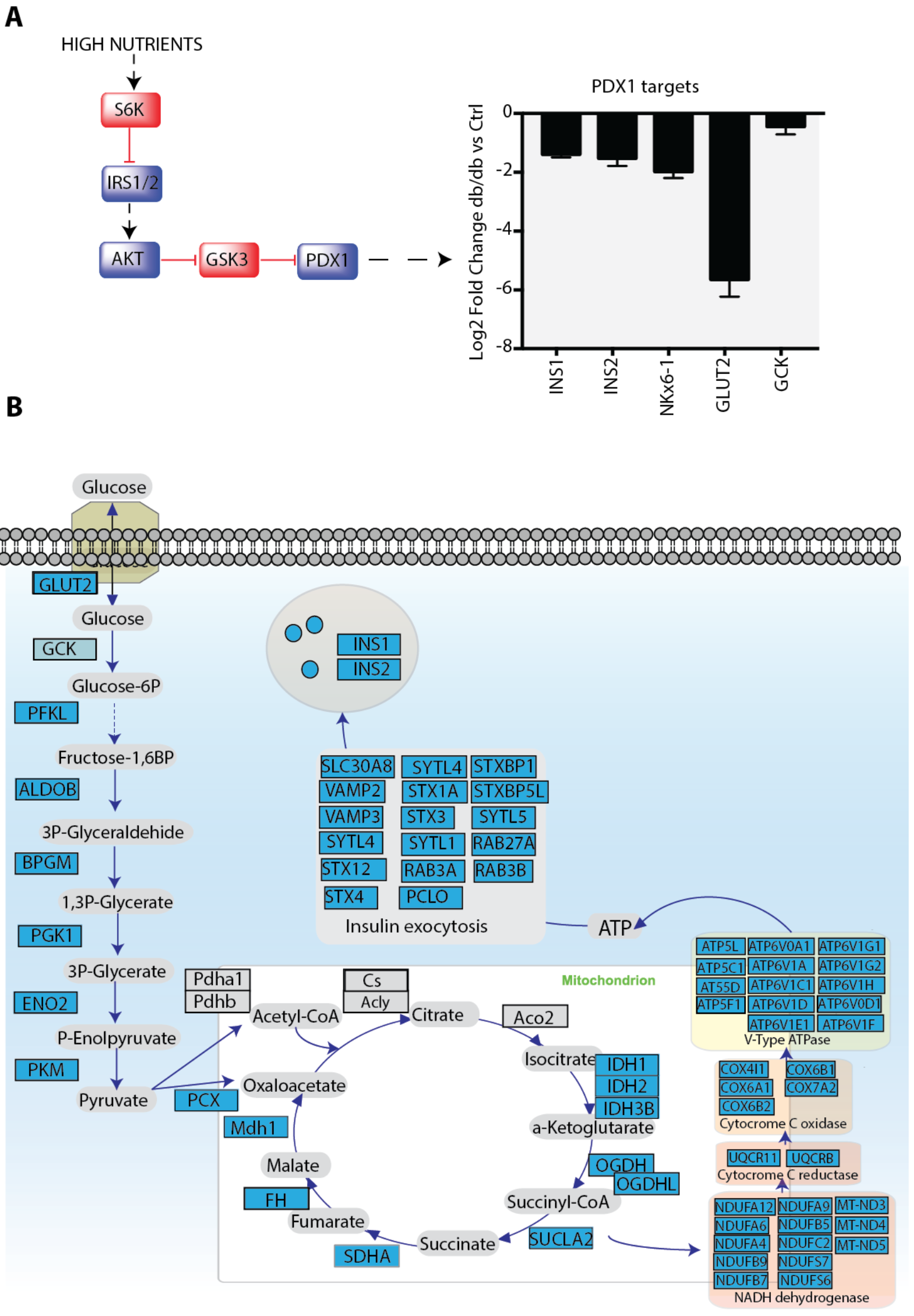
PDX1 down-regulation leads to GLUT2 suppression and glycolysis inhibition. (**A**) Schematic representation of the molecular mechanisms leading to suppression of PDX1 and decreased expression of PDX1 targets (bar graph). (**B**) Schematic representation of glycolysis-TCA-OXPHOS-insulin exocytosis pathways. Down-regulated enzymes are shown as blue squares, while not modulate proteins are grey.

### Glucotoxicity triggers the GSK3-dependent down-regulation of PDX1

During T2D, different factors synergistically contribute to beta cell loss and dysfunction. Chronic or recurrent exposure of beta cells to elevated levels of glucose and lipids (glucolipotoxicity) or to proinflammatory cytokines interfere with beta cell function, contributing to their destruction (Prentki and Nolan, 2006). Given the key role of GSK3 in glucose homeostasis (Humphrey et al., 2010) and having demonstrated that GSK3 inhibits PDX1 and that PDX1 suppression leads to an impairment of insulin secretion machinery in db/db islets, we next aimed to investigate whether glucotoxicity in isolation triggers GSK3 activation in beta cells leading to consequent degradation of PDX1 transcription factor and its targets.

To address this question, we chronically treated isolated INS1e rat beta cells with high glucose and applied our MS-based proteomic and phosphoproteomic workflows to study global changes in these cells (**Fig. 4A**). These experiments enabled the reproducible quantification of around 7,500 proteins and 19,000 phosphosites and their response to chronically elevated glucose levels (**Fig. S9B-C**, **Table S3**). In this experimental system, glucotoxicity drastically affected both the proteome and phosphoproteome, modulating the expression of 4,255 proteins and the phosphorylation level of 5,334 phosphopeptides (Fig?). To investigate and interpret the activation status of the PI3K-AKT-GSK3 and other key signaling pathways, we again integrated our proteomic and phosphoproteomic dataset with the literature-derived signaling network. This revealed that chronic hyperglycaemia in rat beta cells also leads to GSK3 hyperactivation, as specifically demonstrated by six-fold increased phosphorylation of the activating Tyr216 of GSK3. We also observed the subsequent PDX1 phosphorylation by GSK3, causing its proteasomal degradation, as evidenced by its decreased protein levels in our proteomics data (**Fig. 4B**). Consistently, the expression levels of all the identified PDX1 targets, including GLUT2, INS, GCK and PLKR, were also decreased (**Fig. 4C**). Confirming out results in the db/db islets, PDX1 suppression was correlated with suppression of proteins involved in glucose metabolism, TCA cycle, oxidative phosphorylation and insulin secretion (**Fig. 4D**).

**Figure 4.**
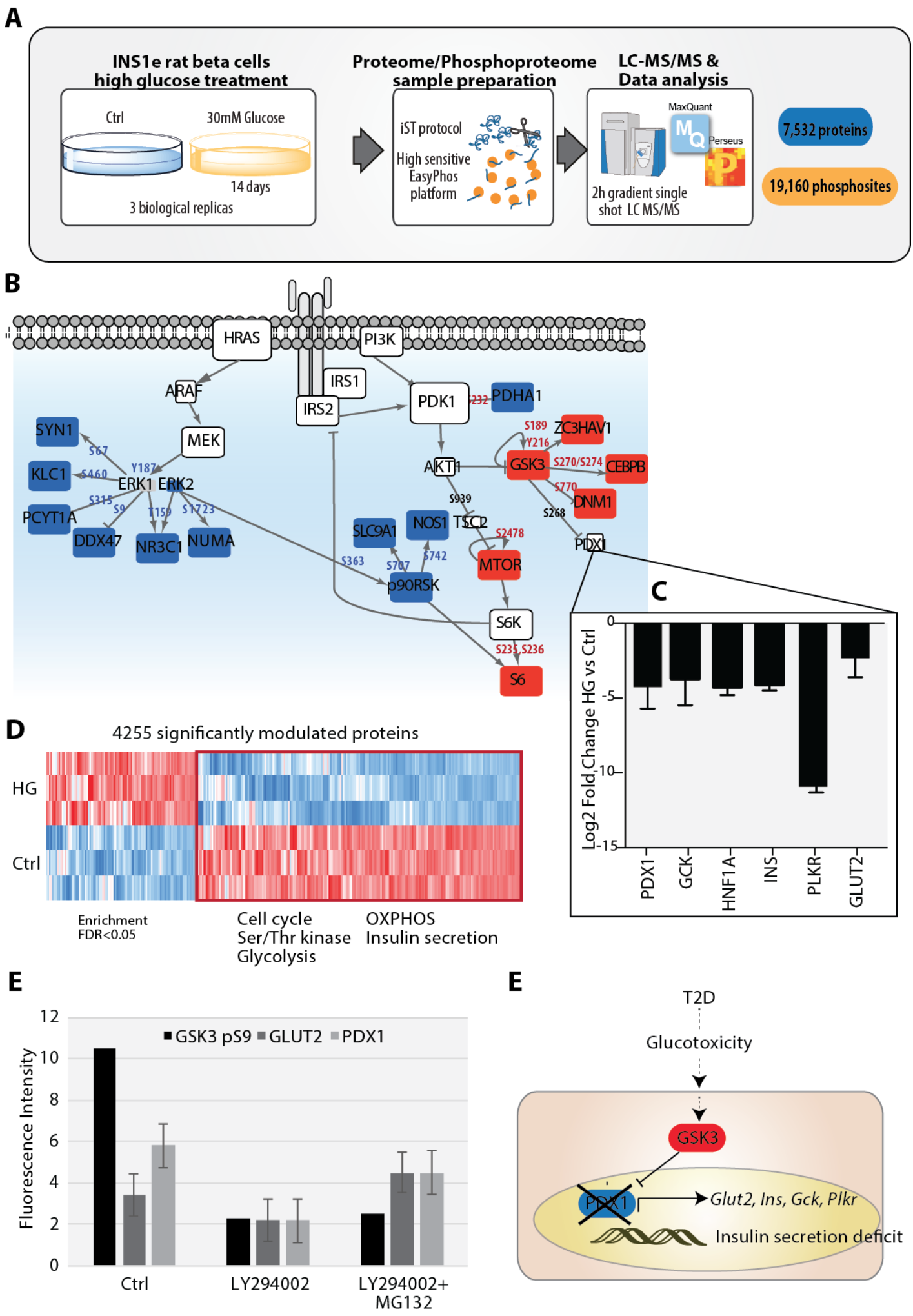
Glucotoxicity triggers GSK3-dependent PDX1 degradation. **A**) Schematic representation of the experimental workflow applied to analyze the proteome and phosphoproteome of INS1e cells after chronic high glucose stimulation. **B**) Proteome and phosphoproteome datasets were overlaid onto a literature-derived signaling network. **C**) Log2 protein expression fold change of PDX1 and its transcriptional targets. **D**) Heatmap of the significantly modulated proteins in INS1e cells after chronic high glucose stimulation. Significantly enriched GO terms and pathways are indicated (FDR<0.07). **E**) Immunofluorescent signal of anti-GSK3 (P), anti-GLUT2 and anti-PDX1. **F**) A proposed model contributing to glucotoxicitydependent beta cell failure

To further assess the functional role of GSK3 in the regulation of PDX1 stability and its transcriptional target GLUT2, we treated human islets with a PI3K inhibitor to trigger GSK3 activation alone or in combination with the proteasome inhibitor MG132. Confirming our hypothesis, GSK3 activation was sufficient to cause PDX1 degradation and GLUT2 suppression (**Fig. 4E, Fig. S10A**). Importantly, GSK3 activation also impaired glucose stimulated insulin secretion (**Fig. S7B**). These effects were completely abrogated by co-treatment with the proteasome inhibitor.

The global proteomics and phosphoproteomic data, together with our functional follow up, thus support a model whereby a diabetic environment, including chronic hyperglycemia and associated glucotoxicity leads to beta cell failure in part through a GSK3-PDX1 dependent axis (**Fig. 4E**).

## Conclusion

Complex diseases are rarely caused by an abnormality in a single gene, but rather by perturbations of global cellular networks. Investigating signaling networks rewiring in diabetic islets has been a major goal of the scientific community, however, the extremely limited amount of material that can be extracted from these structures composed of just a few thousand cells have prevented this so far. Combining a newly developed, highly-sensitive phosphoproteomic workflow with MS-based proteomics we here obtained a first and highly-comprehensive landscape of dynamic changes in protein expression and phosphorylation in obese diabetic mice.

Integrating our large-scale dataset with literature-derived signaling pathways enabled insights into how miss-regulated proteins and signaling events contribute to islets dysfunction. Here we focused on the mechanisms hyperactivation of the key kinase GSK3 in db/db islets, further characterizing its functional role in insulin secretion regulation through a PDX1-dependent mechanism. Tus our proteomic data provide a molecular explanation of the already discovered beneficial effects of GSK3 inhibitor drugs in T2D treatment (MacAulay and Woodgett, 2008; Stein et al., 2011). We hope that in-depth exploration of the resource provided here will aid in the discovery of other potential islet-related diagnostic and therapeutic targets for human T2D.

### Experimental procedure

#### Islet isolation

13 week old C57BLKS-Leprdb homozygous and age-matched C57BLKS-Leprdb/+ heterozygous mice (Charles River Laboratories) were euthanized. The upper abdomen was incised to expose liver and intestines. The pancreas was perfused through the common bile duct with cold collagenase P (from Roche) in saline solution. The pancreas was dissected and placed into a warm collagenase saline solution for 15 min. After enzymatic digestion of the pancreatic tissue, islet were picked and left to recover at 37 °C for 2 hour.

#### Insulin assay

Cells were grown overnight with DMEM low glucose medium, then washed with Krebs-Ringer-Buffer and incubated with starvation buffer for 90 min. Cells were then incubated with high glucose medium (Krebs-Ringer-Buffer supplemented with glucose 16.7 mM and BSA 0.05%) or low glucose medium (Krebs-Ringer-Buffer supplemented with glucose 2.5 mM and BSA 0.05%). Aliquots of the supernatant were assayed for the amount of insulin (insulin assay from Cisbio Bioassays), according to the manufacturer’s protocol.

#### Proteome and phosphoproteome sample preparation

Cells were lysed in SDC lysis buffer containing 4% (w/v) SDC, 100 mM Tris-HCl (pH 8.5). Proteome preparation was done using the in StageTip (iST) method (Kulak et al., 2014). Phosphoproteome preparation was performed as previously described (Humphrey 2015; Humphrey et al., 2018). Per condition, a total of only 200 ug protein input material was lysed, alkylated and reduced in one single step. Then proteins were digested and phosphopeptides enriched by TiO2 beads. After elution, samples were separated by HPLC in a single run (without pre-fractionations) and analyzed by mass spectrometry.

#### Mass spectrometric analyses

The peptides or phosphopeptides were desalted on StageTips and separated on a reverse phase column (packed in-house with 1.8-mm C18-Reprosil-AQ Pur reversed-phase beads) (Dr Maisch GmbH) over 240 min or 270 min (single-run proteome and phosphoproteome analysis). Eluting peptides were electrosprayed and analyzed by tandem mass spectrometry on a Q Exactive HF (Thermo Fischer Scientific) using HCD based fragmentation, which was set to alternate between a full scan followed by up to five fragmentation scans. Proteome and phosphoproteome data were processed and statistically analyzed as described in Supplementary Methods.

#### Cell culture

INS-1E cells (RRID: CVCL_0351) were grown in a humidified atmosphere (5% CO2, 95% air at 37 °C) in monolayer in modified RPMI 1,640 medium supplemented with 10% fetal calf serum, 10 mM Hepes, 100 U ml^−1^, penicillin, 100 mg ml^−1^, streptomycin, 1 mM sodium pyruvate, 50 mM b-mercaptoethanol (all from Gibco) and 0.5% BSA (from Sigma). Human islets were purchased from PELO Biotech and grown according to manufacturer instructions (CellProgen cat n. 35002-04).

## Author contributions

F.S. conceived and M.M. supervised the project. F.S. performed the proteome, phosphoproteome, immunofluorescence and insulin secretion analyses. A.S., F.V. dissected mice, isolated islets. N.K. measured the last batch of proteome samples. S.H.H optimized the workflow for phosphoproteome islets analysis. A.R. helped in the GSK3 activation experiments. F.S. analyzed the data. F.S., N.K, S.H.H, J.G., M.M. wrote the manuscript.

## Acknowledgements

We thank Igor Paron, Korbinian Mayr for technical assistance. These studies were supported by the Max Plank Society for the Advancement of Science.

## Supplementary material

### Supplementary Figures

**Figure S1.**
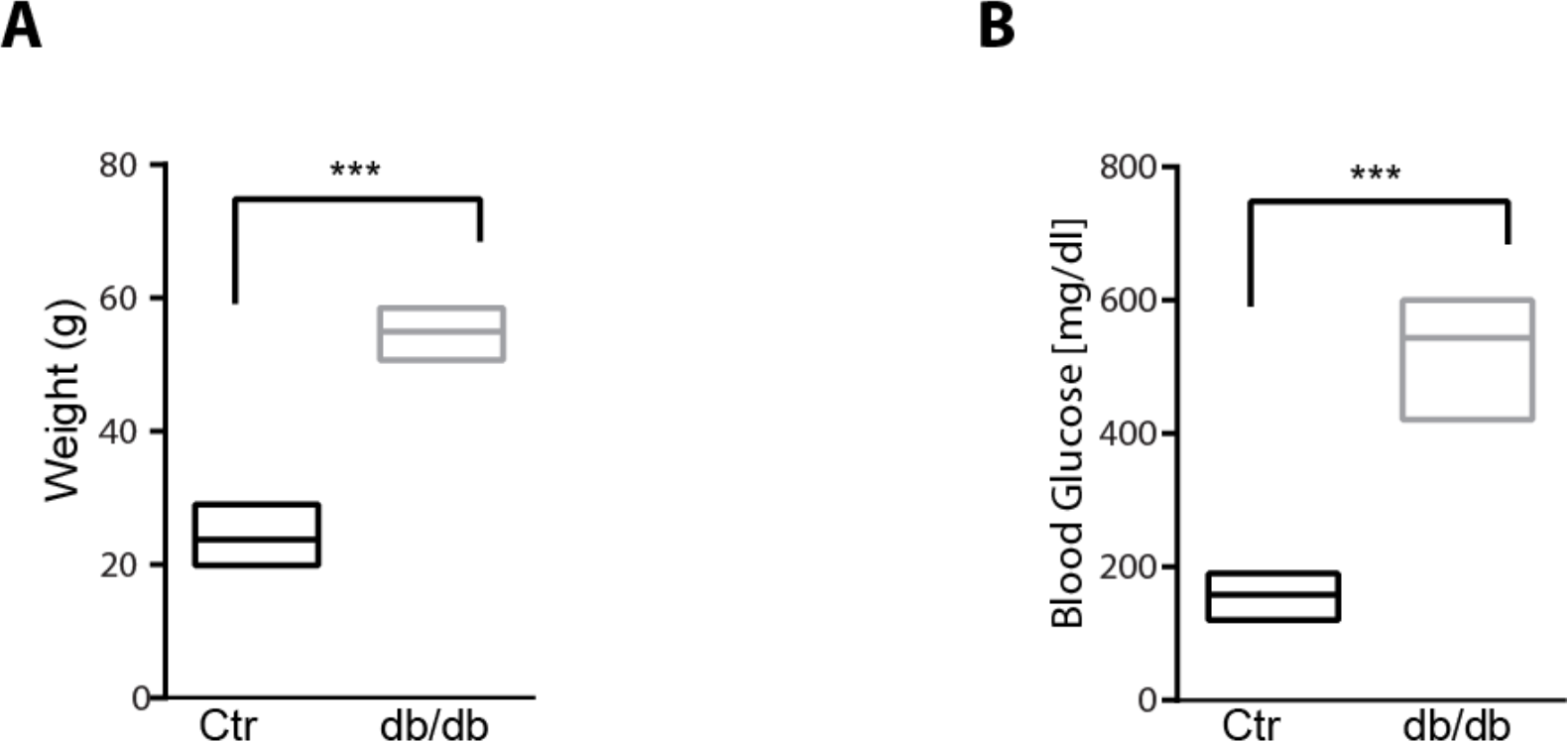
db/db mice are obese with respect to control mice. Body weight (**A**) and plasma glucose (**B**) of db/db and control 13 weeks old mice.

**Figure S2.**
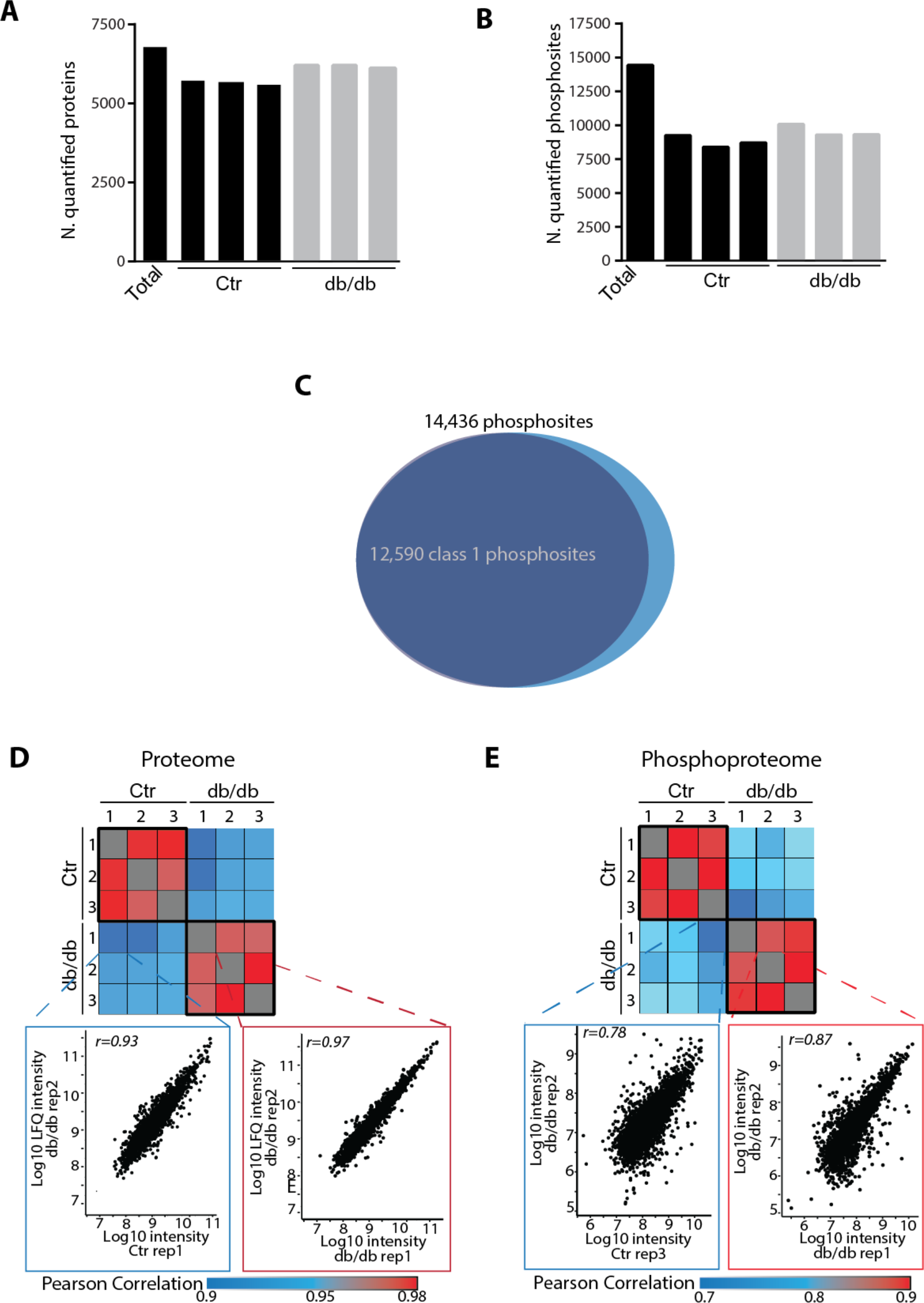
High coverage and reproducibility of proteome and phosphoproteome data. Number of quantified proteins (**A**) and phosphosites (**B**) in biological replicates of different experimental conditions. **C**) Venn diagram showing the proportion of class 1 sites in the whole quantified phosphosites. Heatmap showing the Pearson correlation coefficients between the different biological replicates in the proteome (**D**) and phosphoproteome (**E**) datasets.

**Figure S3.**
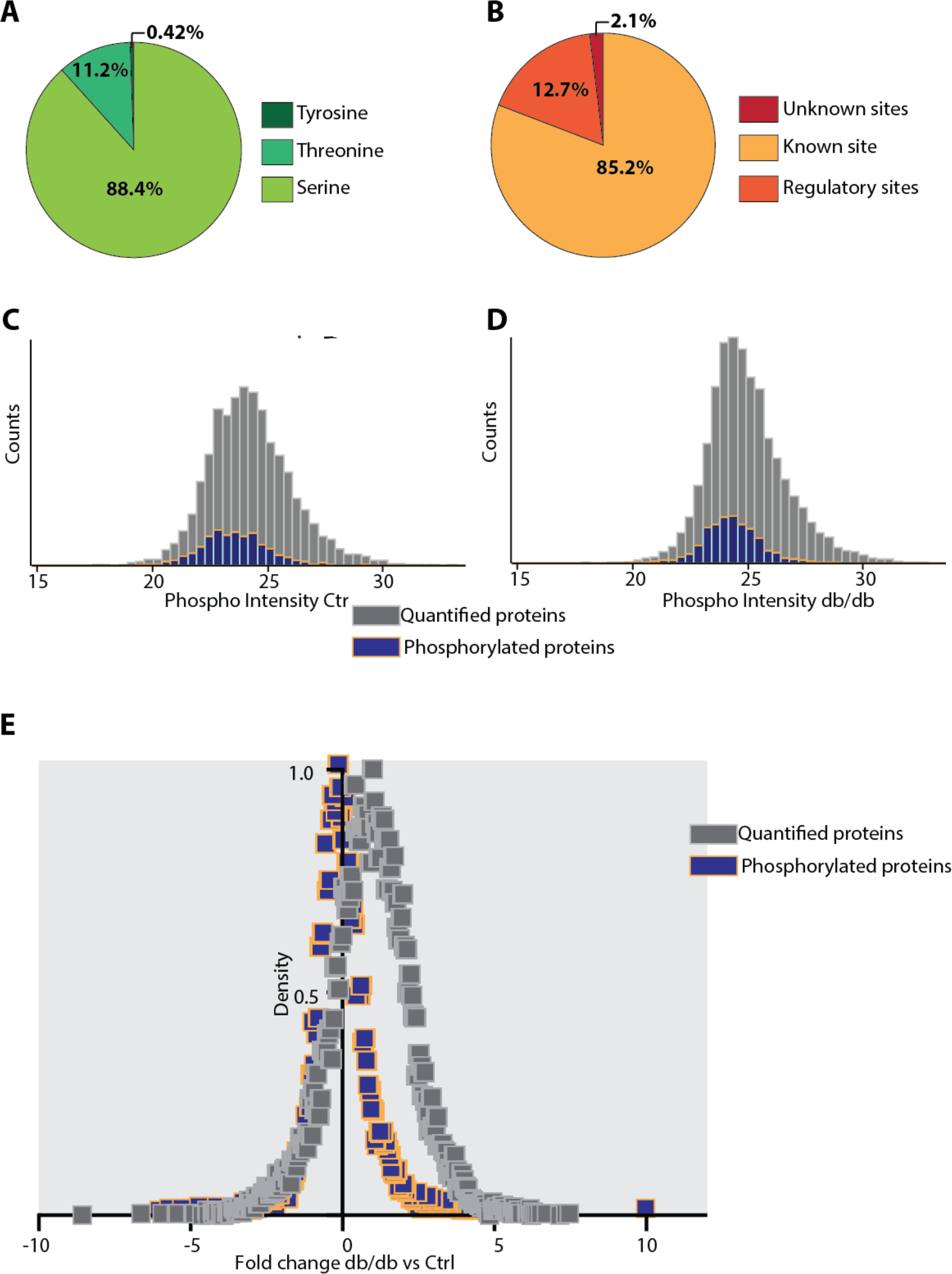
Data summary of the proteomic and phosphoproteomic data. **A**) Distribution of serine, threonine and tyrosine phosphorylation sites. **B**) Phosphosites quantified in our study that are already present in PhosphoSitePlus and annotated as “regulatory sites”. Phosphorylated proteins are distributed in the entire range of measured protein abundances. The distribution of ranked log2 LFQ intensity values in control (**C**) and db/db islets (**D**) are colour coded in grey, while phosphorylation intensity is in blue. **E**) Plot showing the distribution of the amplitudes (fold change of the log2 intensities) calculated for the phosphoproteome (blue) as well as for proteome.

**Figure S4.**
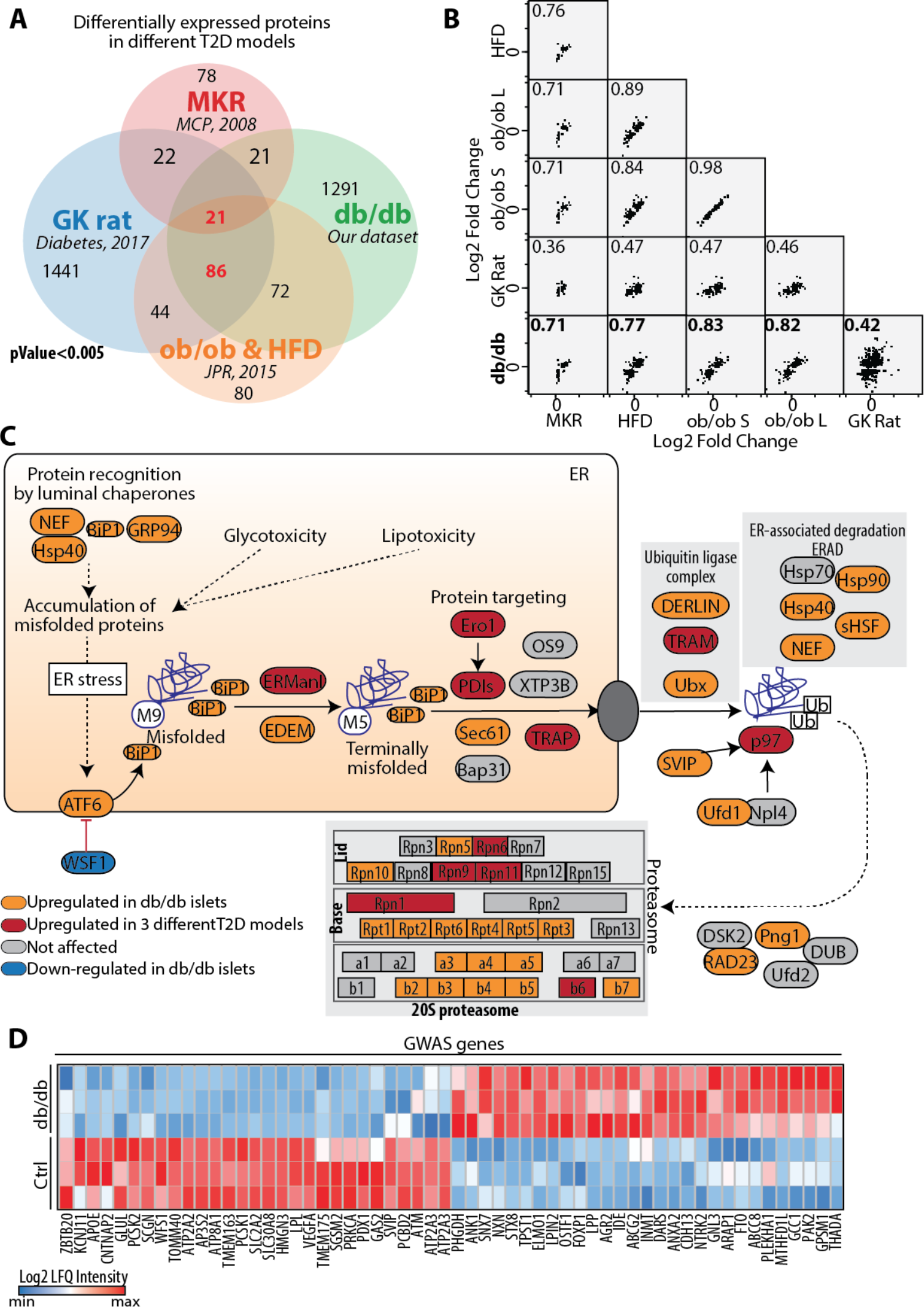
Comparison of proteins significantly modulated in db/db islets, with different T2D literature-derived datasets. **A**) Venn diagram showing the overlap between the differentially expressed proteins derived by our and other proteomic approaches, as indicated. **B**) Multi-scatterplot showing the high correlation between fold changes of T2D-modulated proteins identified by different proteomic approaches. **C**) ER misfolded protein processing and degradation processes are schematically represented. Orange and red proteins are upregulated only in our db/db dataset and in at least 3 different T2D datasets respectively. **D**) Heatmap of protein expression level of genes significantly modulated in db/db islets and found to be significantly associated with T2D by GWAS studies.

**Figure S5.**
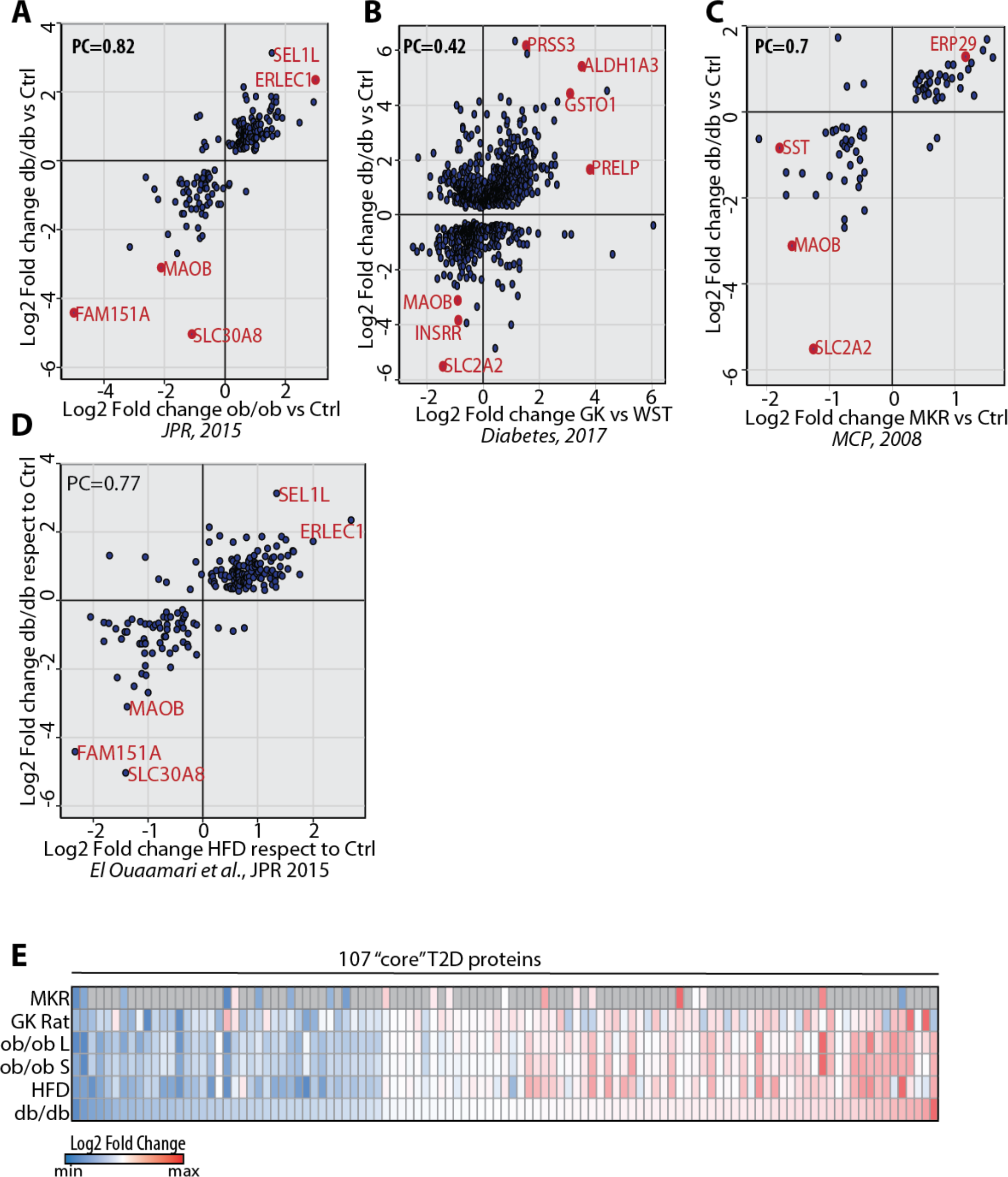
Proteomic comparison of islets derived from different T2D models. **A-B-C-D**) Plots showing the correlation between our dataset and previously published proteomic datasets of islets from the indicated T2D murine and rat models (El Ouaamari et al., 2015; Hou et al., 2017; Lu et al., 2008). **E**) Heatmap of the 107 proteins significantly modulated in islets from at least three different T2D models.

**Figure S6.**
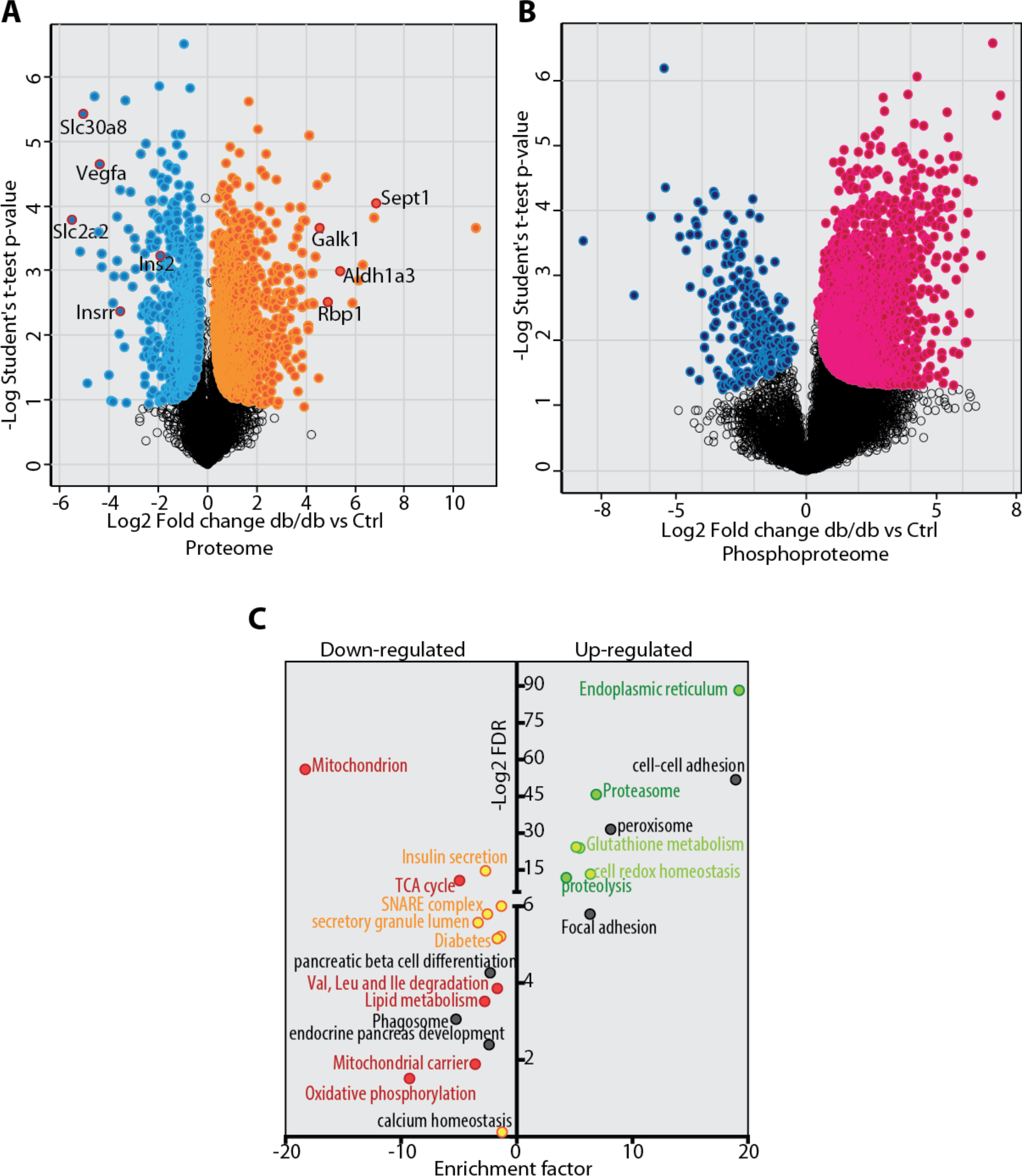
Identification of significantly modulated proteins and phosphosites. Statistically significantly up-and down-regulated proteins (A) and phosphosites (B) were identified by t-test (Benjamin Hochberg FDR < 0.07; S0=0.1) and represented as scatterplot. Each dot represents one protein (A) or one phosphosites (B). **C**) Enrichment and significance of GO-annotations, KEGG pathways and Keywords among the significantly modulated proteins.

**Figure S7.**
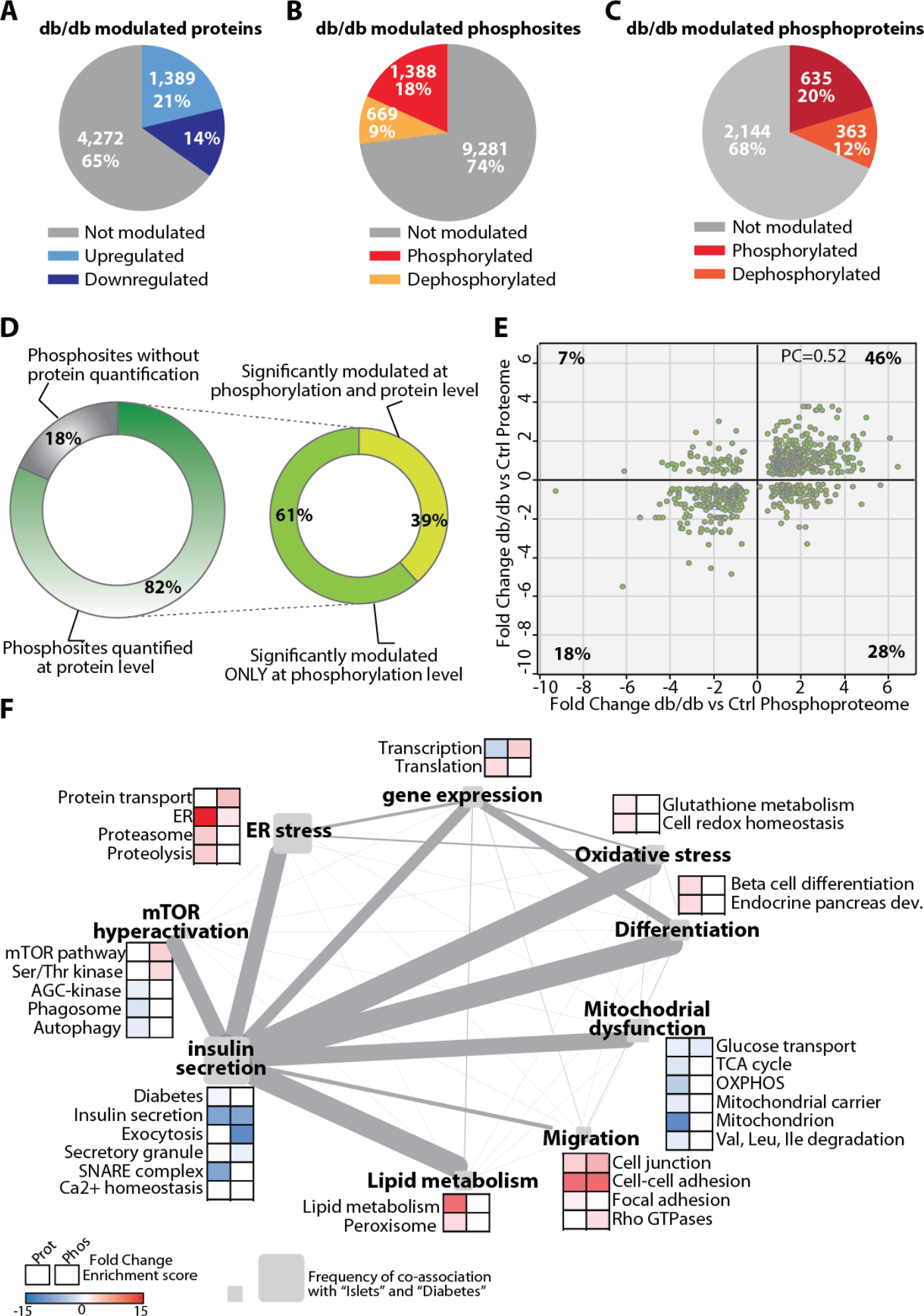
Key biological processes are regulated at the proteome and phosphoproteome levels. **A**) Pie chart indicating the percentage of positively and negatively modulated proteins (**A**), phosphosites (**B**) and phosphoproteins with at least one significantly modulated phosphopeptide (**C**). **D**) Percentage of phosphorylation sites with protein expression level quantified or not (left) and comparison between protein expression and phosphorylation levels for the 82% phosphorylation sites (right). Proteins and phosphorylation sites were considered regulated according to t-test analysis (FDR<0.07; S0=0.1). **E**) Plot showing the comparison of protein expression and phosphorylation level changes for the 39% phosphosites in (D, right). **F**) Clusters of GO term and Kegg pathways enriched in the subset of genes significantly up and down-regulated at the proteome and phosphoproteome level are represented as nodes of a co-citation network. Node size is correlated to the frequency of co-citation of each term with “islets” in literature abstract. Edge thikness is proportial to the “co-citation score”.

**Figure S8.**
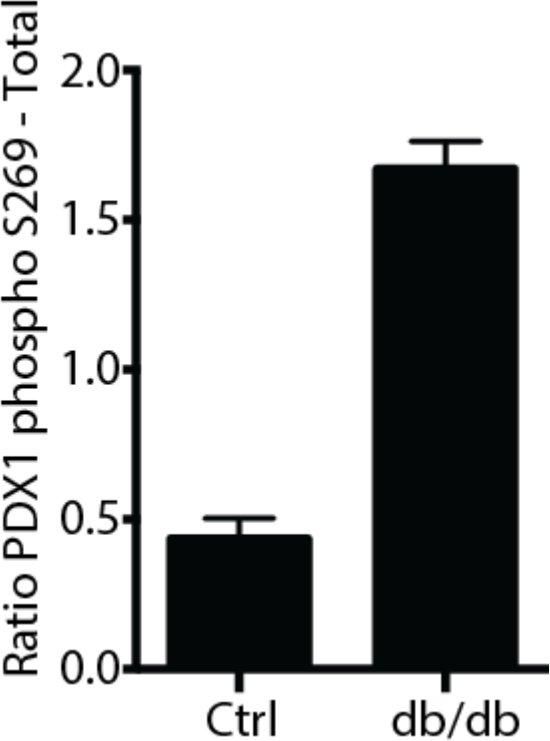
PDX1 phosphorylation in db/db islets. Intensity of the PDX1 phosphopeptide containing S269 (phosphoproteome dataset) normalized to the total protein intensity of the PDX1 protein (LFQ from proteome).

**Figure S9.**
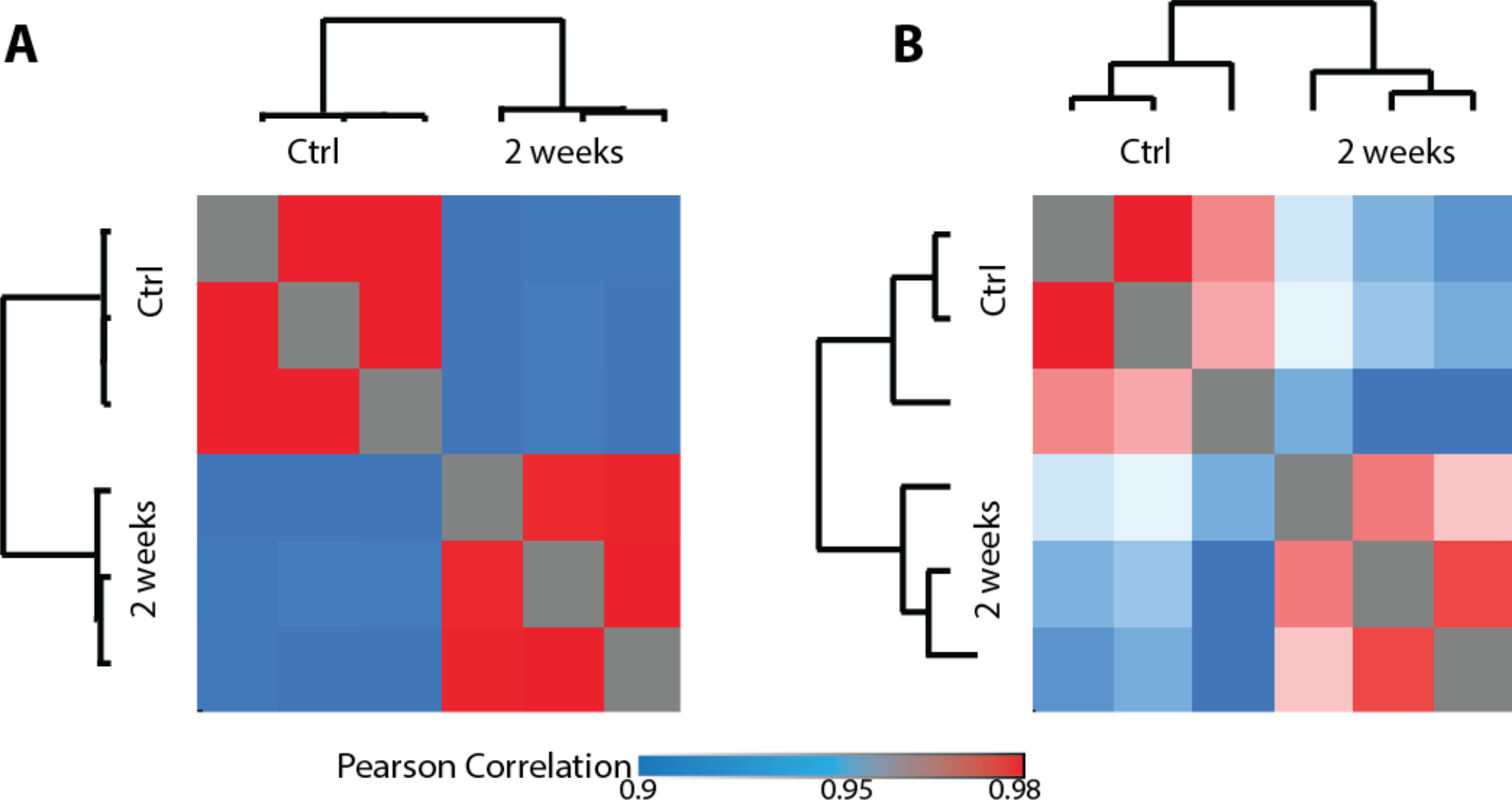
Reproducibility of biological replicates of proteome and phosphoproteome of glucose-treated INS1e cells. Heat map showing the Pearson correlation coefficients between the different biological replicates in the proteome (**B**) and phosphoproteome (**C**) datasets.

**Figure S10.**
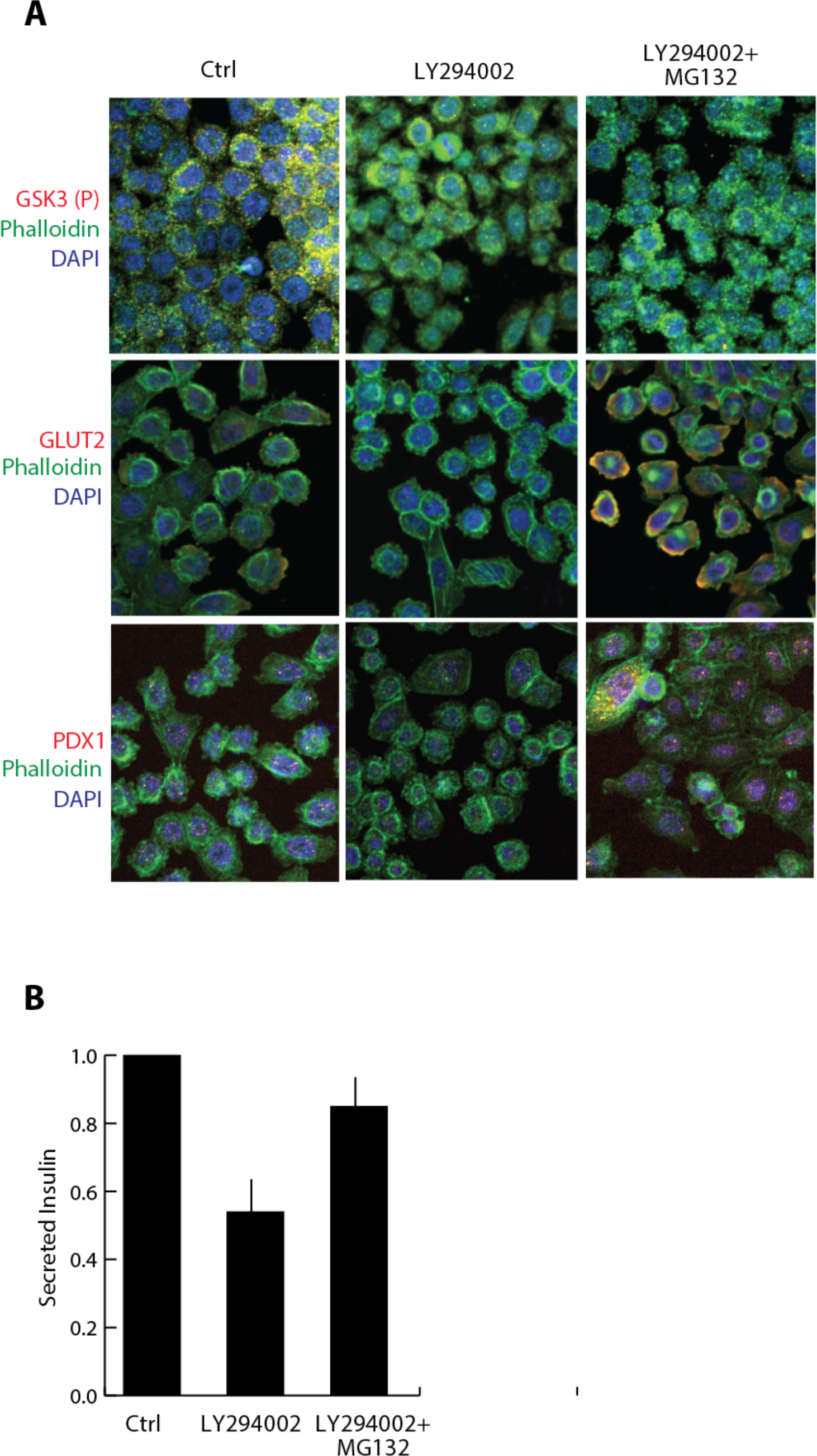
Proteomic comparison of islets derived from different T2D models. **A**) Indirect immunofluorescent analysis of GSK3 S9 (P), GLUT2, PDX1, Phallloidin and DAPI. **B**) Amount of secreted insulin (a.u.) after drug and glucose stimulation measured with an Elisa assay. Median and s.d. of triplicates are shown as a bar graph.

### Supplementary methods

#### Proteome and phosphoproteome data processing

Raw mass spectrometry data were analyzed in the MaxQuant environment (Cox and Mann, 2008), version 1.5.1.6, employing the Andromeda engine for database search. MS/MS spectra were matched against the Mus Murine UniProt FASTA database (September 2014), with an FDR of <1% at the level of proteins, peptides and modifications. Enzyme specificity was set to trypsin, allowing for cleavage N-terminal to proline and between aspartic acid and proline. The search included cysteine carbamidomethylation as a fixed modification, and N-terminal protein acetylation, oxidation of methionine and phosphorylation of serine, threonine tyrosine residue (STY) as variable modifications. Label free proteome analysis was performed in MaxQuant. For proteome and phosphoproteome analysis, where possible, the identity of peptides present but not sequenced in a given run was obtained by transferring identifications across liquid chromatography (LC)-MS runs (‘match between runs’). For phosphopeptide identification, an Andromeda minimum score and minimum delta score threshold of 40 and 17 were used, respectively. Up to three missed cleavages were allowed for protease digestion and peptides had to be fully tryptic.

#### Proteome and phosphoproteome bioinformatics data analysis

Bioinformatic analysis was performed in the Perseus software environment (Tyanova et al., 2016). Statistical analysis of proteome and phosphoproteome was performed on logarithmized intensities for those values that were found to be quantified in any experimental condition. Phosphoproteome intensities were normalized by subtracting the median intensity of each sample. To identify significantly modulated proteins and phosphopetides across two different conditions, we performed a Student t-Test with a permutation-based FDR cutoff of 0.07 and S0=0.1. Categorical annotation was added in Perseus in the form of GO biological process (GOBP), molecular function (GOMF), and cellular component (GOCC), KEGG pathways and kinase substrate motifs (extracted from HPRD). Concerning the kinase substrate motifs, we performed a 1D annotation enrichment analyses to identify statistically significant enriched kinase-substrates motifs in db/db islets (Cox and Mann, 2012). Multiple hypothesis testing was controlled by using a Benjamini-Hochberg FDR threshold of 0.05. Then for each kinase-substrate motif the corresponding pValue and score are assigned. While a score near 1 indicates a positive enrichment, a score near −1 means a negative enrichment of the category.

#### Combining proteome and phosphoproteome data with a prior knowledge signaling network

This strategy has been previously developed and applied by our group (Sacco et al., 2012; Sacco et al., 2016). Kinase-substrate relationships were extracted by PhosphositePlus (Kinase-substrate dataset) and SIGNOR (Hornbeck et al., 2015; Perfetto et al., 2016) and were mapped onto the complete human proteome. Then the network was first filtered to maintain only relationships between proteins that were identified in our proteomic analysis of db/db islets or INS1e cells. This network was used as a scaffold to overlay the changes at the phosphoproteome level. Next the network was filtered according to the following rules: i) “leaf nodes” (connected only by one edge) whose phosphorylation was not quantified in our phospho-proteomics approach were excluded; ii) “Leaf nodes” whose phosphorylation was not modulated in db/db islets or hyperglycaemic INS1e cells were also excluded and iii) those residues (edges) whose phosphorylation status was not supported by our experimental data were eliminated. This filtering procedure yielded a much simpler network that was easier to analyse. For those proteins having multiple regulatory residues and whose phosphorylation was strongly changed in opposite direction, we considered only those sites quantified in a singly phosphorylated peptide (multiplicity =1).

#### Immunofluorescence microscopy

Human pancreatic islets were treated with LY294002 with or without MG132. Cells were fixed for 10 min in 4% paraformaldehyde (EM Sciences), washed with PBS, 0.5% Triton X-100, and permeabilized with blocking solution (0.1% Triton, 10% fetal calf serum) for 30 min. Cells were incubated, in blocking solution, with anti-GSK3 (P) (RRID:AB_10013750), anti-GLUT2, anti-PDX1 and anti FITC phalloidin (RRID:AB_2315147) for 1 h at room temperature. Cells were rinsed in PBS and incubated with secondary antibody for 30 minutes. Cells were stained with 4’,6-diamidino-2-phenylindole in PBS, 0.1% Triton for 5 min at room temperature. Cells were analyzed by indirect immunofluorescence microscopy.

